# Comparing the Citation Performance of PNAS Papers by Submission Track

**DOI:** 10.1101/036616

**Authors:** Philip M. Davis

## Abstract

**Purpose:** To determine whether papers contributed by National Academy of Sciences (NAS) members perform differently than direct submissions.

**Data/Methods:** 55,889 original papers published in *PNAS* from 1997 through 2014. Regression analysis measuring total citations, controlling for editorial track (Contributed, Direct, Communicated), date of publication, and paper topic.

**Main findings:** Contributed papers consistently underperformed against Direct submissions, receiving 9% fewer citations, *ceteris paribus*. The effect was greatest for Social Sciences papers (12% fewer citations). Nonetheless, the main effect has attenuated over the past decade, from 13.6% fewer citations in 2005 to just 2.2% fewer citations in 2014.

**Significance:** Successive editorial policies placing limits, restrictions, and other qualifications on the publication privileges of NAS members may be responsible for the submission of better performing Contributed papers.

## Background

Over the years, successive editorial policies for the *Proceedings of the National Academy of Sciences (PNAS)* have placed limits, restrictions, and other qualifications on the publication privileges of NAS members. The intentions of these policy changes were to improve upon the transparency of the editorial process and external review, and secondly, to encourage the submission of higher-quality papers. Should these policies have their intended effects, we should expect to observe a gradual improvement in the performance of papers contributed by NAS authors compared to direct submissions.

*PNAS* currently allows two manuscript submission tracks^1^: *Direct*, in which papers are subject to single-blind peer review; and *Contributed*, in which a NAS member may secure their own reviewers and submit their comments alongside the paper. A third submission track (Communicated), in which papers are sent directly to a NAS member, who is responsible for overseeing the peer review process, was abolished in 2010.^2^

### Example changes to PNAS publication policies

1972 – NAS members can Contribute or Communicate up to 10 papers per year.

1989–1992. Quota reduced to 6 papers/yr.

1993–1995. Quota reduced to 5 papers/yr.

1994 – July. Conflict of Interest (COI) disclaimer is added to NAS members’ paper.

1995 – Dec. COI disclaimer requirement is removed.

1996 – Quota reduced to 4 papers/yr.

2001 – Names of reviewers are required when Contributing or Communicating papers.

2006 – Jan. NAS member must be a corresponding author on Contributed papers.

2006 – Aug. PNAS begins listing author contributions on papers.

2007 – May. Two reviewers are required for Contributed papers

2008 – April. NAS members with a significant COI with their Contributed papers are required to use Direct submission track.

2010 – July. PNAS eliminates Communicated submissions.

2014 – COI decision of 2008 repealed.

2014 – April. Voluntarily listing reviewer names on contributed papers.

2015 – Oct. Requiring reviewer names on contributed papers.

## Data and methods

The dataset includes 55,889 original research papers published in *PNAS* from 1997 through 2014, years for which submission track information was available from the publisher. Perspectives, Commentary, and other non-research papers were excluded from the dataset. For each *PNAS* paper, the complete citation record was extracted from the *Web of Science* (Thomson Reuters) during the first week of November 2015 and matched with the *submission track* (Direct, Contributed, Communicated) and journal *section* (e.g. Biological Sciences—Microbiology) using the paper’s Digital Object Identifier (DOI) or by matching the paper’s volume and page number.

Our primary objective was to evaluate the performance of papers by submission track. To arrive at a relative performance measure, we compared the performance of Contributed and Communicated papers against Direct Submission papers for each year of publication. Performance was measured by the total number of citations to a *PNAS* paper. As citation distributions are highly skewed, we normalized the distribution by adding one citation to each paper before taking the natural logarithm of the result.^3^

Our primary analysis involved building a linear regression model, using log(*total citations*) as the response variable, and *submission track* (Contributed, Communicated) as two indicator variables. We controlled for publication date within each year by using the journal’s *issue* number. Finally, we controlled for the topic of each paper by using both the *primary domain* of each paper (Biological Sciences, Physical Sciences, Social Sciences), as well as its specific *field* (e.g. Microbiology). As some fields are classified under multiple domains—for example, Anthropology papers are classified under both Biological Science and Social Sciences—we nested the *field* variable within the *domain* variable.

## Results

*PNAS* grew by 44% over the observation period, from 2,437 research papers published in 1997 to 3,521 papers in 2014 (Table 1), during which, the proportion of Direct submissions increased from one-in-five (22%) *PNAS* papers to over three-quarters (77%). In contrast, Contributed submissions have remained relatively constant. In 2010, *PNAS* eliminated the Communicated submission track.

**Table 1.** *PNAS* papers by submission track, 1997—2014.

| Year | Submission Track |  |  | Total |
| --- | --- | --- | --- | --- |
|  | Communicated | Contributed | Direct |  |
| 1997 | 1203 (49%) | 709 (29%) | 525 (22%) | 2437 |
| 1998 | 1227 (47%) | 701 (27%) | 691 (26%) | 2619 |
| 1999 | 1128 (45%) | 680 (27%) | 679 (27%) | 2487 |
| 2000 | 863 (36%) | 701 (29%) | 841 (35%) | 2405 |
| 2001 | 715 (29%) | 755 (31%) | 989 (40%) | 2459 |
| 2002 | 836 (30%) | 767 (27%) | 1201 (43%) | 2804 |
| 2003 | 816 (31%) | 742 (28%) | 1081 (41%) | 2639 |
| 2004 | 784 (30%) | 767 (29%) | 1060 (41%) | 2611 |
| 2005 | 774 (25%) | 887 (28%) | 1498 (47%) | 3159 |
| 2006 | 796 (24%) | 831 (25%) | 1655 (50%) | 3282 |
| 2007 | 746 (22%) | 775 (22%) | 1937 (56%) | 3458 |
| 2008 | 597 (17%) | 795 (23%) | 2076 (60%) | 3468 |
| 2009 | 492 (13%) | 764 (21%) | 2461 (66%) | 3717 |
| 2010 | 256 (7%) | 754 (20%) | 2701 (73%) | 3711 |
| 2011 | . | 773 (22%) | 2754 (78%) | 3527 |
| 2012 | . | 786 (21%) | 2954 (79%) | 3740 |
| 2013 | . | 872 (23%) | 2973 (77%) | 3845 |
| 2014 | . | 807 (23%) | 2714 (77%) | 3521 |
| Total | 11233 (20%) | 13866 (25%) | 30790 (55%) | 55889 |

In every year, from 1997 to 2014, Contributed papers underperformed against Direct submissions (Figure 1, Table 2). Controlling for the date of publication and journal section (field of study), Contributed papers received 9% fewer citations, on average, than Direct submissions. The effect was greatest for Social Sciences papers (12% fewer citations). Nonetheless, the main effect has attenuated over the past decade, from 13.6% fewer citations in 2005 to just 2.2% fewer citations in 2014.

**Figure 1.**
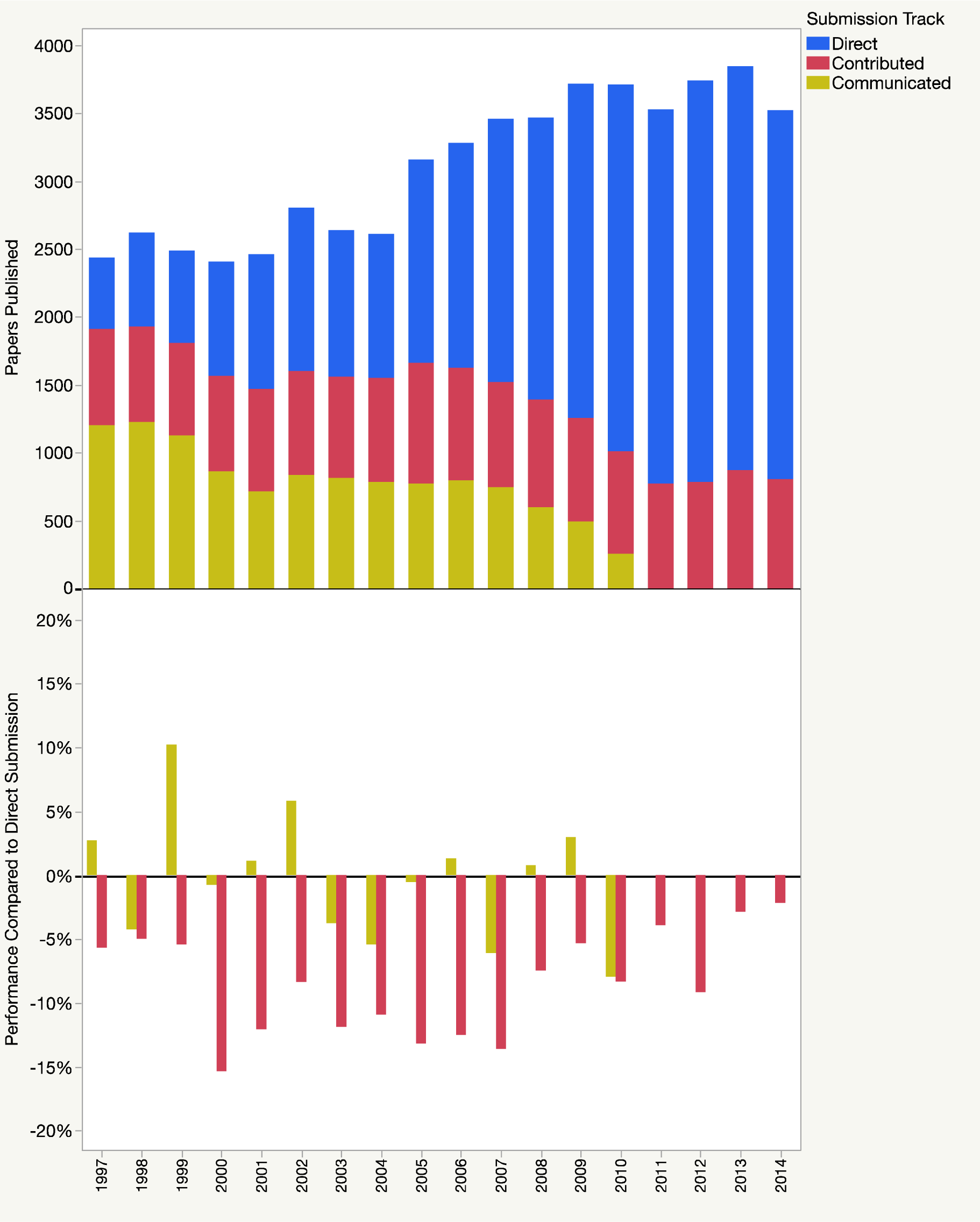
Research papers published by track (top) and citation performance compared to Direct submissions (bottom). Publication and performance numbers are found in Tables 1 and 2.

**Table 2.**
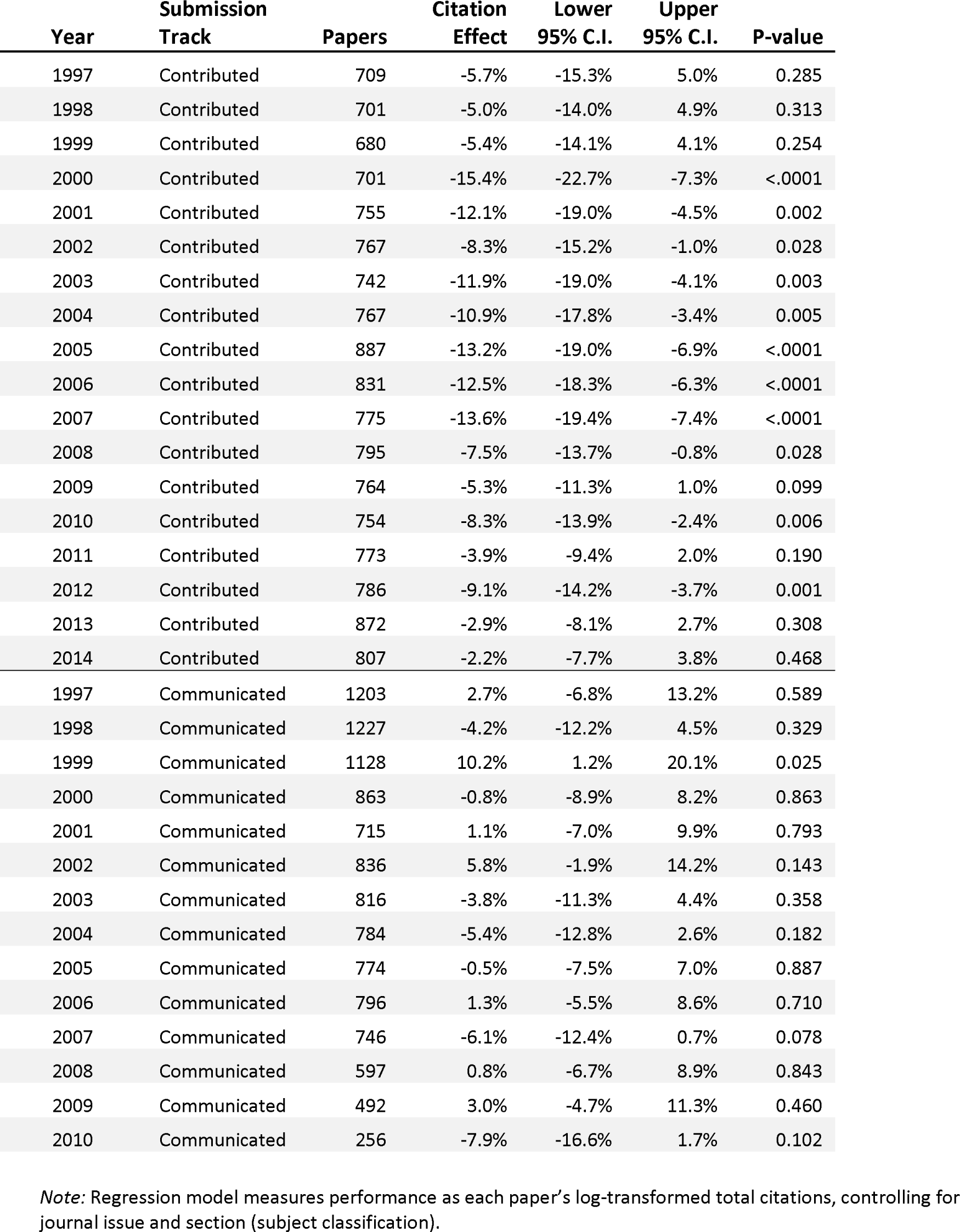
Performance of *PNAS* research papers compared to Direct submissions by year of publication.

| Year | Submission Track | Papers | Citation Effect | Lower 95% C.I. | Upper 95% C.I. | P-value |
| --- | --- | --- | --- | --- | --- | --- |
| 1997 | Contributed | 709 | -5.7% | -15.3% | 5.0% | 0.285 |
| 1998 | Contributed | 701 | -5.0% | -14.0% | 4.9% | 0.313 |
| 1999 | Contributed | 680 | -5.4% | -14.1% | 4.1% | 0.254 |
| 2000 | Contributed | 701 | -15.4% | -22.7% | -7.3% | <.0001 |
| 2001 | Contributed | 755 | -12.1% | -19.0% | -4.5% | 0.002 |
| 2002 | Contributed | 767 | -8.3% | -15.2% | -1.0% | 0.028 |
| 2003 | Contributed | 742 | -11.9% | -19.0% | -4.1% | 0.003 |
| 2004 | Contributed | 767 | -10.9% | -17.8% | -3.4% | 0.005 |
| 2005 | Contributed | 887 | -13.2% | -19.0% | -6.9% | <.0001 |
| 2006 | Contributed | 831 | -12.5% | -18.3% | -6.3% | <.0001 |
| 2007 | Contributed | 775 | -13.6% | -19.4% | -7.4% | <.0001 |
| 2008 | Contributed | 795 | -7.5% | -13.7% | -0.8% | 0.028 |
| 2009 | Contributed | 764 | -5.3% | -11.3% | 1.0% | 0.099 |
| 2010 | Contributed | 754 | -8.3% | -13.9% | -2.4% | 0.006 |
| 2011 | Contributed | 773 | -3.9% | -9.4% | 2.0% | 0.190 |
| 2012 | Contributed | 786 | -9.1% | -14.2% | -3.7% | 0.001 |
| 2013 | Contributed | 872 | -2.9% | -8.1% | 2.7% | 0.308 |
| 2014 | Contributed | 807 | -2.2% | -7.7% | 3.8% | 0.468 |
| 1997 | Communicated | 1203 | 2.7% | -6.8% | 13.2% | 0.589 |
| 1998 | Communicated | 1227 | -4.2% | -12.2% | 4.5% | 0.329 |
| 1999 | Communicated | 1128 | 10.2% | 1.2% | 20.1% | 0.025 |
| 2000 | Communicated | 863 | -0.8% | -8.9% | 8.2% | 0.863 |
| 2001 | Communicated | 715 | 1.1% | -7.0% | 9.9% | 0.793 |
| 2002 | Communicated | 836 | 5.8% | -1.9% | 14.2% | 0.143 |
| 2003 | Communicated | 816 | -3.8% | -11.3% | 4.4% | 0.358 |
| 2004 | Communicated | 784 | -5.4% | -12.8% | 2.6% | 0.182 |
| 2005 | Communicated | 774 | -0.5% | -7.5% | 7.0% | 0.887 |
| 2006 | Communicated | 796 | 1.3% | -5.5% | 8.6% | 0.710 |
| 2007 | Communicated | 746 | -6.1% | -12.4% | 0.7% | 0.078 |
| 2008 | Communicated | 597 | 0.8% | -6.7% | 8.9% | 0.843 |
| 2009 | Communicated | 492 | 3.0% | -4.7% | 11.3% | 0.460 |
| 2010 | Communicated | 256 | -7.9% | -16.6% | 1.7% | 0.102 |

*Note:* Regression model measures performance as each paper's log-transformed total citations, controlling for journal issue and section (subject classification).

For Communicated papers, the results were mixed. In half of the years, Communicated papers performed better than Direct Submissions; in the other half, they performed worse.

Inaugural papers (typically the first Contributed paper published by a NAS member) received 8% more citations (95% C.I. 4% to 12%, p=0.0001) than other Contributed papers and performed statistically no different than Direct submissions (citation deviance = +2%, 95% C.I. -1% to 6%, p=0.170).

After their second calendar year of publication, 3.8% (526 of 13866) of Contributed papers remained uncited, compared to 2.6% (789 of 30790) of Direct submissions. Uncitedness varied by *PNAS* field (Figure 2, Table 3). Controlling for date of publication and journal section, Contributed papers were nearly twice as likely to remain uncited (Odds Ratio = 1.7, 95% C.I. 1.5 to 1.9, Chi^2^<.0001). Uncitedness among Contributed papers, however, has been declining at 3% per annum (95% C.I. -5% to -1%, p=0.01).

**Figure 2.**
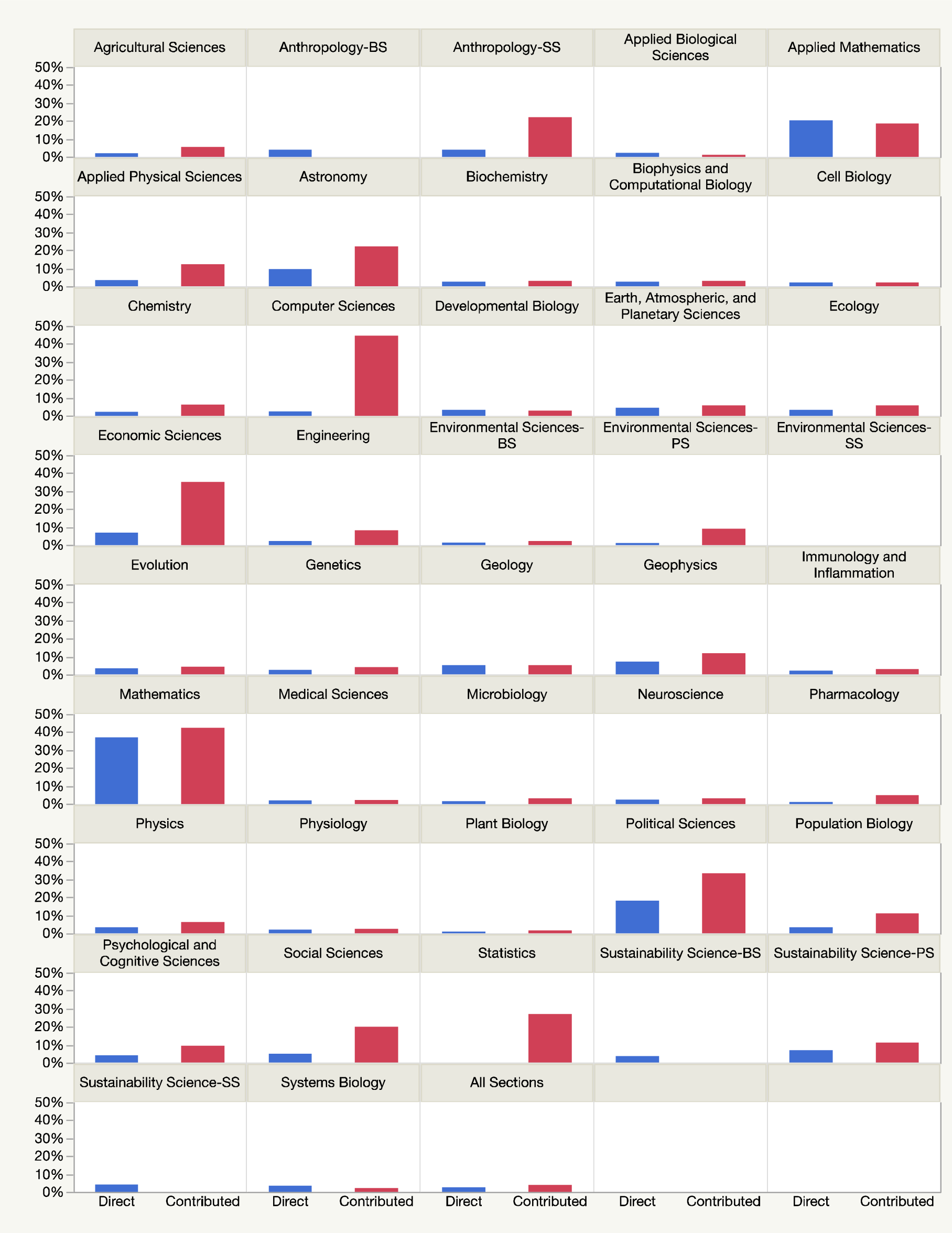
Percentage of uncited papers by section in their first two calendar years of publication (Blue=Direct, Red=Contributed). Numbers and percentages are found in Table 3.

**Table 3.** Uncited *PNAS* papers in their first two calendar years of publication.

| Section | DIRECT |  |  | CONTRIBUTED |  |  |
| --- | --- | --- | --- | --- | --- | --- |
|  | N (zero) | N (all) | % zero | N (zero) | N (all) | % zero |
| Agricultural Sciences | 3 | 156 | 2% | 2 | 36 | 6% |
| Anthropology-BS | 7 | 181 | 4% | 0 | 55 | 0% |
| Anthropology-SS | 5 | 130 | 4% | 13 | 59 | 22% |
| Applied Biological Sciences | 10 | 471 | 2% | 2 | 177 | 1% |
| Applied Mathematics | 21 | 103 | 20% | 13 | 70 | 19% |
| Applied Physical Sciences | 19 | 554 | 3% | 19 | 156 | 12% |
| Astronomy | 4 | 42 | 10% | 4 | 18 | 22% |
| Biochemistry | 87 | 3663 | 2% | 62 | 2105 | 3% |
| Biophysics and Computational Biology | 60 | 2520 | 2% | 25 | 863 | 3% |
| Cell Biology | 44 | 2133 | 2% | 24 | 1134 | 2% |
| Chemistry | 25 | 1197 | 2% | 31 | 502 | 6% |
| Computer Sciences | 1 | 42 | 2% | 4 | 9 | 44% |
| Developmental Biology | 30 | 951 | 3% | 12 | 453 | 3% |
| Earth, Atmospheric, and Planetary Sciences | 12 | 277 | 4% | 2 | 36 | 6% |
| Ecology | 25 | 803 | 3% | 9 | 162 | 6% |
| Economic Sciences | 6 | 89 | 7% | 13 | 37 | 35% |
| Engineering | 5 | 219 | 2% | 4 | 49 | 8% |
| Environmental Sciences-BS | 3 | 252 | 1% | 1 | 47 | 2% |
| Environmental Sciences-PS | 2 | 201 | 1% | 4 | 44 | 9% |
| Environmental Sciences-SS | 0 | 22 | 0% | 0 | 4 | 0% |
| Evolution | 49 | 1442 | 3% | 22 | 506 | 4% |
| Genetics | 36 | 1438 | 3% | 36 | 875 | 4% |
| Geology | 7 | 135 | 5% | 3 | 58 | 5% |
| Geophysics | 7 | 97 | 7% | 6 | 51 | 12% |
| Immunology and Inflammation | 36 | 1753 | 2% | 33 | 1104 | 3% |
| Mathematics | 46 | 124 | 37% | 14 | 33 | 42% |
| Medical Sciences | 46 | 2590 | 2% | 35 | 1594 | 2% |
| Microbiology | 27 | 2021 | 1% | 20 | 656 | 3% |
| Neuroscience | 75 | 3339 | 2% | 47 | 1599 | 3% |
| Pharmacology | 4 | 380 | 1% | 8 | 164 | 5% |
| Physics | 12 | 368 | 3% | 13 | 213 | 6% |
| Physiology | 15 | 731 | 2% | 7 | 303 | 2% |
| Plant Biology | 10 | 1103 | 1% | 6 | 386 | 2% |
| Political Sciences | 2 | 11 | 18% | 1 | 3 | 33% |
| Population Biology | 4 | 125 | 3% | 2 | 18 | 11% |
| Psychological and Cognitive Sciences | 23 | 569 | 4% | 13 | 140 | 9% |
| Social Sciences | 5 | 102 | 5% | 7 | 35 | 20% |
| Statistics | 0 | 62 | 0% | 7 | 26 | 27% |
| Sustainability Science-BS | 4 | 114 | 4% | 0 | 11 | 0% |
| Sustainability Science-PS | 3 | 44 | 7% | 1 | 9 | 11% |
| Sustainability Science-SS | 4 | 96 | 4% | 0 | 19 | 0% |
| Systems Biology | 5 | 140 | 4% | 1 | 47 | 2% |
| All Sections | 789 | 30790 | 2.6% | 526 | 13866 | 3.8% |

Notes: BS-Biological Sciences; PS-Physical Sciences; SS-Social Sciences

In spite of their general underperformance, the 10% most-cited Contributed papers *outperformed* the top 10% of Direct submissions, receiving 8% more citations, on average (95% C.I. 5% to 11%, p=<.0001) over the entire dataset (Figure 3, Table 4). Consistent with our main findings, these effects have also been attenuating over the past decade and were not detectable in the last five years of publication (Table 4).

**Figure 3.**
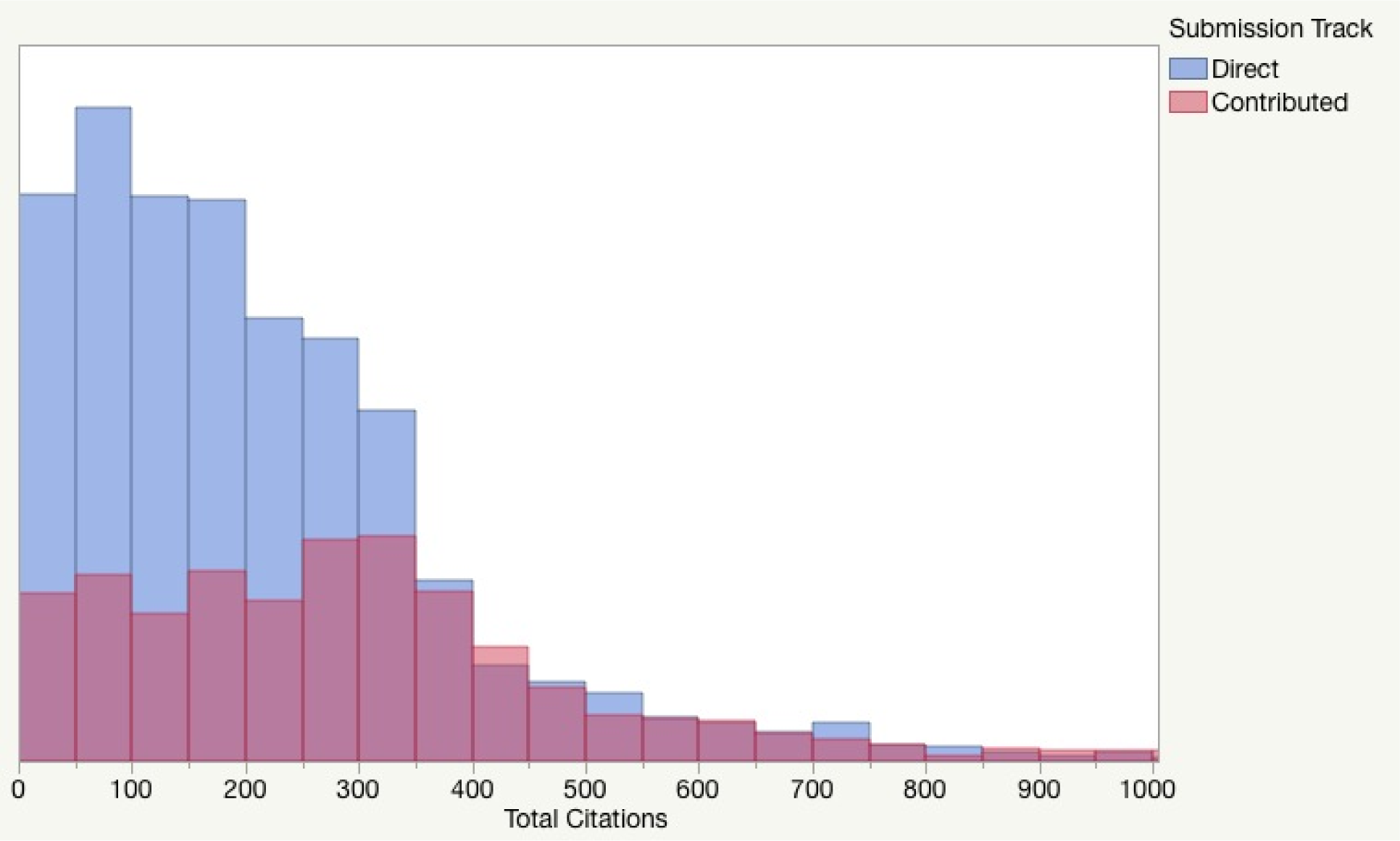
Histogram of the 10% most-cited *PNAS* papers for each year (1997-2014) by submission track. Regression results for each year of publication are found in Table 4.

**Table 4.**
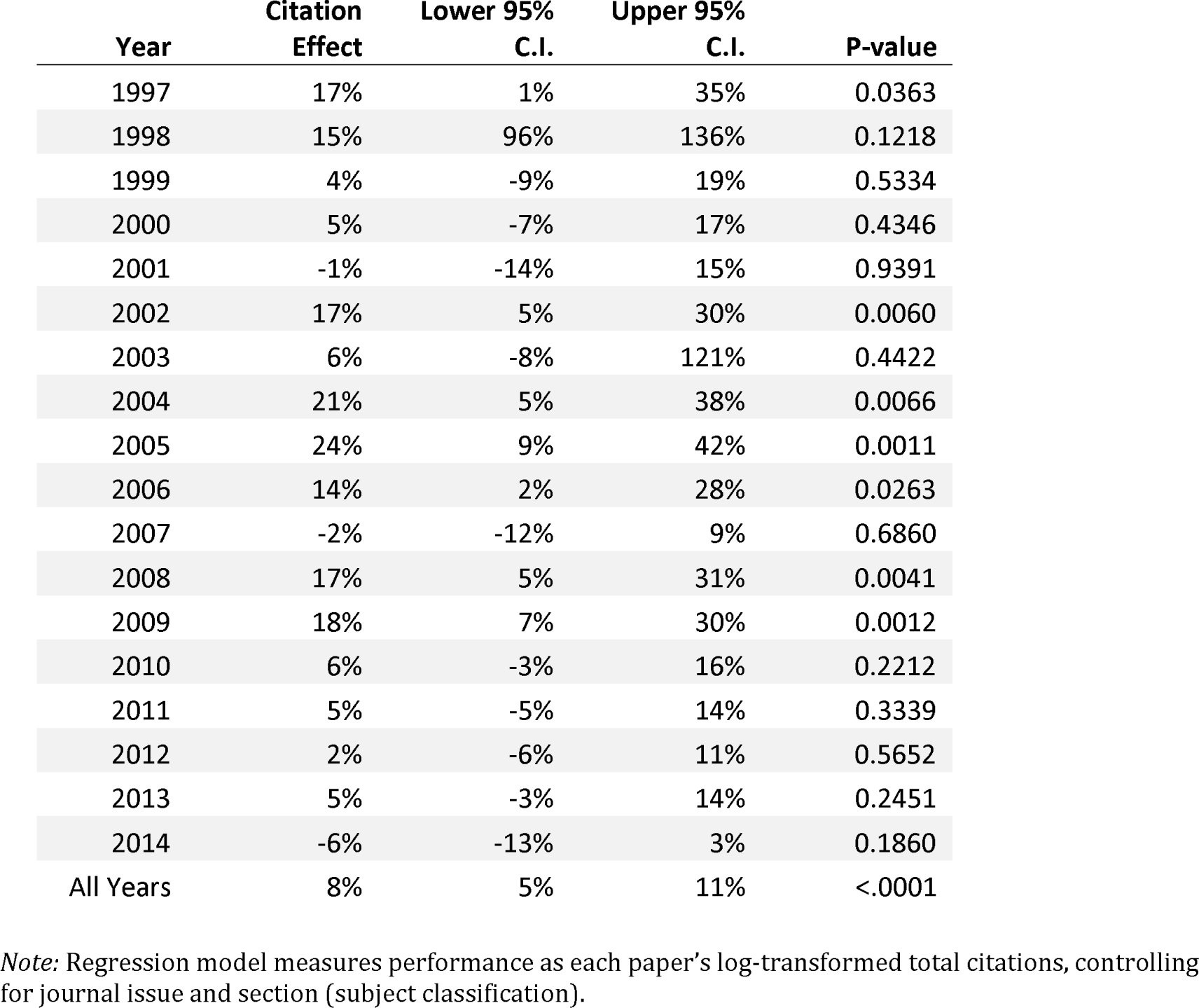
Citation performance of the 10% most-cited Contributed *PNAS* papers compared to Direct submissions.

| Year | Citation Effect | Lower 95% C.I. | Upper 95% C.I. | P-value |
| --- | --- | --- | --- | --- |
| 1997 | 17% | 1% | 35% | 0.0363 |
| 1998 | 15% | 96% | 136% | 0.1218 |
| 1999 | 4% | -9% | 19% | 0.5334 |
| 2000 | 5% | -7% | 17% | 0.4346 |
| 2001 | -1% | -14% | 15% | 0.9391 |
| 2002 | 17% | 5% | 30% | 0.0060 |
| 2003 | 6% | -8% | 121% | 0.4422 |
| 2004 | 21% | 5% | 38% | 0.0066 |
| 2005 | 24% | 9% | 42% | 0.0011 |
| 2006 | 14% | 2% | 28% | 0.0263 |
| 2007 | -2% | -12% | 9% | 0.6860 |
| 2008 | 17% | 5% | 31% | 0.0041 |
| 2009 | 18% | 7% | 30% | 0.0012 |
| 2010 | 6% | -3% | 16% | 0.2212 |
| 2011 | 5% | -5% | 14% | 0.3339 |
| 2012 | 2% | -6% | 11% | 0.5652 |
| 2013 | 5% | -3% | 14% | 0.2451 |
| 2014 | -6% | -13% | 3% | 0.1860 |
| All Years | 8% | 5% | 11% | <.0001 |

*Note:* Regression model measures performance as each paper's log-transformed total citations, controlling for journal issue and section (subject classification).

## Prior Work

Studying *PNAS* papers published between June 1, 2004 and April 26, 2005 and using a similar regression model, Rand and Pfeiffer^4^ reported that Contributed papers received 10% fewer citations compared to Direct submissions, a result that is consistent with our findings for those years (see Table 2). Similarly, they reported that the 10% most-cited Contributed papers outperformed the 10% most-cited Direct submissions, consistent with our findings for 2004 and 2005 (see Table 4). Aldhous^5^ reported that that only a very small number of NAS members used the Contributed track frequently; the majority of members published less than one contributed paper per year, on average.

## Study Limitations

This study compared the performance of papers by *submission track* not by NAS *member status*. While all Contributed papers must include at least one NAS author, it is not known whether Direct submissions included any NAS authors. Having the membership status of each author would have permitted us to disentangle author effects from submission track effects.

## Conclusions

The difference in citation performance between Contributed and Direct submissions has been declining over the past decade, suggesting that successive editorial policies intended to improve upon the transparency and submission of higher-quality contributed papers is the primary explanation.

## Footnotes

1 PNAS Editorial Policies. http://www.pnas.org/site/authors/guidelines.xhtml

2 Schekman, R. PNAS will eliminate Communicated submissions in July 2010. http://dx.doi.org/10.1073/pnas.0909515106

3 Adding one citation to each paper allows for papers that have not yet been cited to be included in the analysis as the log of zero is a logical impossibility. While this procedure right-shifts the distribution, our analysis focuses on the *comparative* performance of tracks and not their absolute values.

4 Rand DG & Pfeiffer T (2009) Systematic Differences in Impact across Publication Tracks at PNAS. *PLoS ONE* 4(12):e8092. http://dx.doi.org/10.1371/journal.pone.0008092

5 Aldhous P (2014) Scientific publishing: The inside track. *Nature* 510:330-332. http://dx.doi.org/10.1038/510330a

